# SpecHap: a diploid phasing algorithm based on spectral graph theory

**DOI:** 10.1101/870972

**Authors:** Yonghan Yu, Lingxi Chen, Xinyao Miao, Shuai Cheng Li

**Affiliations:** Department of Computer Science, City University of Hong Kong. Tat Chee Avenue 83, Kowloon, Hong Kong, 999077, China

## Abstract

Haplotype phasing is essential to study diploid eukaryotic organisms. High-throughput sequencing, including next-generation sequencing and third-generation sequencing from different technologies, brings possibilities for haplotype assembly. Although there exist multiple haplotype phasing algorithms, only a few are portable across sequencing technologies with the premise of efficiency and accuracy. Herein, we proposed SpecHap, a novel haplotype assembly tool that leverages spectral graph theory, transforming haplotype phasing into an algebraic problem. On both in silico and whole-genome-sequencing datasets, SpecHap consumed less memory and required less CPU time, yet achieved comparable accuracy comparing to state-of-art methods across all the test instances of next-generation sequencing, linked-reads, high-throughput chromosome conformation capture sequencing, PacBio single-molecule real-time sequencing and Oxford Nanopore long-reads sequencing data. Furthermore, SpecHap successfully phased an individual *Ambystoma mexicanumm*, a species with gigantic diploid genomes, within 6 CPU hours and 945MB peak memory usage, while other tools failed to yield results either due to a memory overflow (40GB) or a time limit excess (5 days). Our results demonstrated that SpecHap is scalable, efficient and accurate for diploid phasing, supporting diverse sequencing platforms.

## INTRODUCTION

Human and many other animals possess diploid genomes with paternal and maternal sets of chromosomes (Zheng *et al*., 2016). The two haplotypes, sequential presentation of heterogeneous single nucleotide variation (SNV), small insertion and deletion (INDEL), large genome rearrangement through structure variation or copy number variation, depict the most genetic variation of diploid eukaryotes (Glusman *et al*., 2014). Phasing, reconstruction of specific allele sequence on individual chromosome, is the fundamental for understanding compound heterozygosity (Tewhey *et al*., 2011). Many studies have addressed the importance of haplotype phasing, including but not limited to allelic differential, epigenomic regulations, and population development (Consortium *et al*., 2012, 2010, Onuchic *et al*., 2018, Tan *et al*., 2018). Furthermore, several studies have proclaimed that haplotype analysis led to disease susceptibility, which is concealed in unphased single nucleotide polymorphism (SNP) signals (Begnini *et al*., 2010, Conrad *et al*., 2006, Musone *et al*., 2008, Trégouët *et al*., 2009).

With advances in high-throughput sequencing, multiple sequencing protocols enable credible identifications and linkages of genetic variants, greatly contributing towards the construction of single individual haplotype (SIH). Next-generation sequencing (NGS) technologies, including the Ion Torrent S5 system (Life Technologies), have been widely used to study the proximal haplotypes leveraging paired-end reads (Qi *et al*., 2018). For high-throughput chromosome conformation capture (Hi-C), HaploSeq verified 95% of heterozygous variants could be phased correctly (Panconesi and Sozio, 2004, Selvaraj *et al*., 2013). Further, transinteractions between homologous chromosomes complicate the process of phasing. Moreover, segmental duplication and simple repeats are likely related to incorrect contig orientation based on Hi-C data (Burton *et al*., 2013, Kaplan and Dekker, 2013, Tan *et al*., 2018). With higher NGS throughput and cost-effectiveness, 10x synthetic long reads (SLRs) protocol provides barcoded linked-reads (*>*100kb long-range information) and is suitable for assembling and phasing (Zhang *et al*., 2019, Zheng *et al*., 2016). Despite the high error rate of ∼15%, the third-generation sequencing (TGS), including single-molecule real-time (SMRT) sequencing technology from Pacific Biosciences (PacBio) and Oxford Nanopore Technologies (ONT), offer ultra-long reads with moderate coverage, significantly promoting the resolution of haplotypes (Edge *et al*., 2017, Pollard *et al*., 2018). 20 **************

Several algorithms exist to assemble haplotypes from sequencing reads on diverse protocols. These methods could be summarized into three categories: optimization by reads (fragments) partitioning, by minimum error correction (MEC) and by haplotype likelihood. Several routines are adopted to solve the optimization including dynamic programming, Markov chain Monte Carlo (MCMC), and heuristic graph cut. **(i)** FastHare, a fast algorithm based on fragments partitioning optimization (Panconesi and Sozio, 2004) produces significant-high error rate (Edge *et al*., 2017). **(ii)** Bansal *et al*. proposed an MCMC-based algorithm HASH and a max-cut heuristic algorithm HapCUT (Bansal and Bafna, 2008) that both assembling haplotype by minimising the MEC. DCHap, a divide-and-conquer algorithm that combines fragment partitioning and MEC optimization, designed for TGS data, achieved a comparable phasing accuracy (Li and Lin, 2020). **(iii)** Based on graph max-cut algorithm, RefHap presented a formulation for SIH (Duitama *et al*., 2010). As a general algorithm for human haplotype assembly, HapCUT2, adopts max-cut computations in the haplotype graph to find the haplotype with maximum likelihood (Edge *et al*., 2017). However, computationally efficient, scalable and accurate methods are still demanded with the advance of sequencing technology.

Herein, we developed SpecHap, a novel fast and accurate scalable algorithm for diploid haplotype phasing designed for multiplex sequencing platforms, especially for TGS error-prone long-reads. Instead of iteratively approaching haplotype by optimizing MEC or haplotype likelihood (Edge *et al*., 2017), SpecHap assembled haplotype efficiently by transforming haplotype phasing into a standard linear algebra problem with spectral graph theory. We benchmarked SpecHap with four state-of-art phasing software and demonstrated its comparable accuracy on diverse sequencing protocol. Moreover, SpecHap phased an individual of amphibian specie *Ambystoma mexicanumm* (Axolotl), which possesses one of the largest genomes (32 billion base pairs), with ∼30× PacBio SMRT long-reads (Nowoshilow *et al*., 2018, Weisrock *et al*., 2018).

## MATERIALS AND METHODS

SIH involves assembling of fragment which contains allele information at heterozygous variants loci inferred by sequence alignment to reference genome. The heterozygous variants may be determined by the same set of individual genome re- sequencing data. In a simplified situation where the individual genome differs from the reference only by SNVs and the fragments are error-free, haplotype might be obtained through partitioning the fragments into two groups in which fragments contain no conflicting alleles. The elementary bi-partitioning approach is unsuitable to assemble haplotype in real cases where reads might be error-prone.

Several measurements are introduced to handle high throughput error-prone reads. One of the widely adopted criteria, MEC refers to the number of alleles within fragments for each to be consistent with estimated haplotype strain. Most haplotype assembling strategies focus on optimizing MEC objective function, while several algorithms adopt a probabilistic model for haplotype assembly (Duitama *et al*., 2012, Edge *et al*., 2017). These methods are algorithms that iteratively infer better haplotype. In this study, we proposed SpecHap, a novel algorithm that assembles haplotype with spectral graph theory.

### Spectral Graph Theory

A simple bi-partitioning elaboration of spectral graph theory could be described as follows. Assume an undirected graph *G* consists of *N* vertices *V* ={*v*_1_,*v*_2_,…,*v*_*N*_}, and *M* edges *E* ={*e*_1_,*e*_2_,…,*e*_*M*_} where each edge *e*_*m*_ =(*v*_*i*_,*v*_*j*_),*i* ≠ *j*. The adjacency matrix *A*^*N ×N*^ is constructed for graph *G* to store the linkage relationship between a pair of vertices *v*_*i*_, *v*_*j*_, such that *A*_*i,j*_ = *w*_*i,j*_ if (*v*_*i*_, *v*_*j*_) ∈ *E* and *A*_*i,j*_ = 0, otherwise. Assume *d*_*i*_ is the degree of a given vertex *v*_*i*_ in graph *G*. The degree matrix *D* is a diagonal matrix with *i*-th diagonal element assigned to *d*_*i*_: *D*_*i,j*_ = *d*_*i*_ if *i* = *j, D*_*i,j*_ = 0 otherwise. Then, by definition, the unnormalised Laplacian matrix *L* of graph *G* is constructed as *L* = *D*−*A*. According to spectral graph theory, these *N* vertices can be grouped into two clusters by the element sign (+,−) of the eigenvector corresponding to the second smallest eigenvalue of *L* (Fiedler vector) (Chung and Graham, 1997). The result obtained through spectral analysis often outperforms classic clustering algorithm and can be solved efficiently through linear algebra (Von Luxburg, 2007).

By constructing a graph with each vertex representing an allele at a heterozygous variant locus, an edge between two vertices of different variants loci demonstrates a potential pair-wise haplotype with edge weight stands for the score or level of confidence for the corresponding haplotype. With the linkage graph structure, spectral graph analysis can be applied and haplotype string is deduced through the Fiedler vector.

#### Linkage graph construction

We first extract the fragments information from the alignment and variant file using modified extractHAIRS (Edge *et al*., 2017). The fragments are then disassembled into pair-wise linkage information, which could be summarized into the weight of corresponding edge. The average genomic span between pair-wise linked heterozygous variants loci varies among different sequencing technologies. For whole-genome sequencing (WGS) with short insertion, reads are scanned for het-variants loci in a local region with hundreds of base pairs length; With TGS reads, the distance between two connected variant might be extended up to millions of base pair; For 10x linked-reads, barcodes are also examined with its range inferred from the alignment result and hence the linked fragment may cover variants loci separated by thousands of base pairs; For Hi-C data, linkages among het-variants loci apart from millions of base pairs might be extracted.

With the fragment information, a *linkage graph* is constructed. In the linkage graph, a pair of nodes for each heterozygous variant locus represents its two parallel alleles (**Fig1:a,b,c**). For variant locus *S*_*i*_, we denote its two parallel alleles as *S*_*iu*_and *S*_*iū*_respectively. Similarly, for the other het-SNV locus *S*_*j*_, we have *S*_*jv*_ and 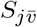. Each fragment is then disassembled into pair-wise linkage between alleles of corresponding variants (*S*_*i*_ and *S*_*j*_). There are two types of linkages: direct and indirect. Taking alleles *S*_*iu*_ and *S*_*jv*_ as an instance, the direct linkage is provided by the fragment covering *S*_*iu*_ and *S*_*jv*_, while the indirect linkage is provided by the fragment covering parallel alleles *S*_*iū*_ and 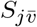. We integrate both direct and indirect linkages into an edge connecting *S*_*iu*_ and *S*_*jv*_nodes. The linkage graph can also be generalized in situations where each node representing a phased haplotype block, enabling SpecHap to connect/extend already phased haplotype.

**Figure 1.**
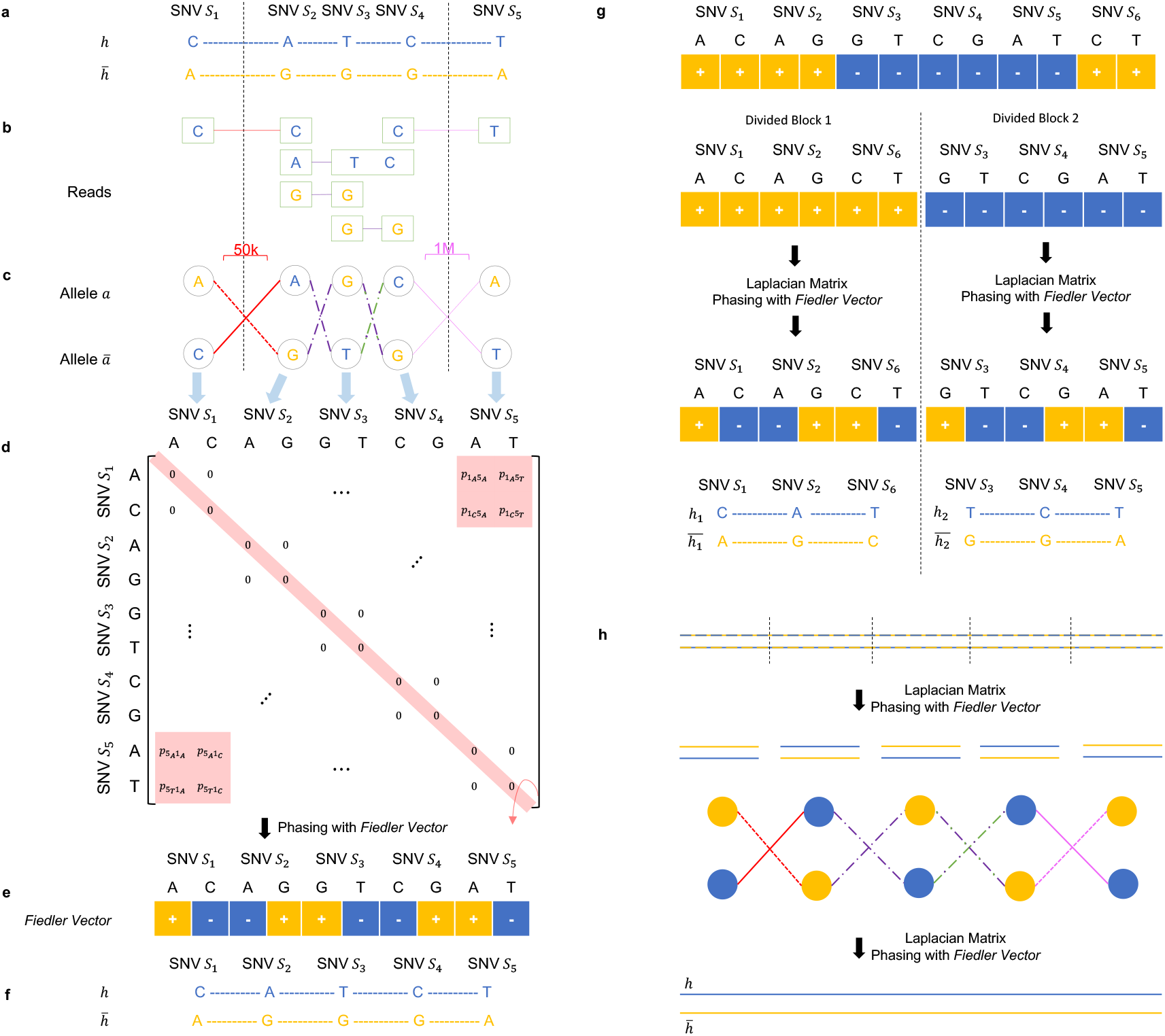
Illustration of SpecHap Algorithm. **(a)** Real haplotypes to be resolved. Blue and yellow alleles are allied to haplotypes *h* and 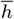 respectively. **(b)** Observed sequencing reads. Green frame refers to single-end read; red line implies linkage supported by 10x barcode; purple line represents the WGS paired-end linkage; pink line stands for the Hi-C or TGS linkage. **(c)** The linkage graph. The solid edge refers to the direct linkage from sequencing reads, the dashed edge is the indirect linkage from parallel alleles pair, and the “solid-dashed” alternation represents edges composed from direct and indirect linkage together. **(d)** The weighted adjacent matrix for graph. e.g. *p*_5*A*_1_*C*_ and *p*_1*C*_ 5_*A*_ are the same, they denote the linkage probability between SNV *S*_5_ allele “A” and SNV *S*_1_ allele “C”. **(e)** Fiedler vector procured by spectral graph theory. **(f)** Resolved haplotypes *h* and 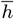 determined by Fiedler vector. **(g)** Simple example demonstrating haplotype construction from Fiedler Vector that guides variants partitioning. **(h)** Connecting haplotype blocks from phased sub-blocks by re-applying Spectral Graph Theory.

#### Logarithmic likelihood as edge weight

To calculate the edge weight of linkage graph, we introduce a likelihood heuristic method based on the Phred probability of variant in fragments. Consider *q*[*j*] as the likelihood that variant *j* is incorrect on fragment *R*_*i*_. Given any haplotype string *h*, the likelihood of observing fragment *R*_*i*_with *j* variants is deduced as:

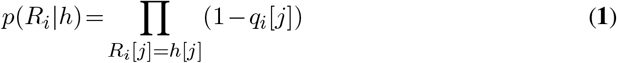

Given a self-complemented haplotype pair 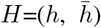, the likelihood *p*(*R*_*i*_|*H*) that a fragment *R*_*i*_ been observed is generalized as 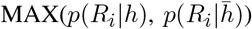). The likelihood that fragment set *R* been observed is:

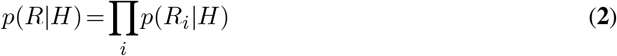

Our linkage graph incorporates edges that representing conflicting haplotype. To decrease the noise and computational load, we define the edge weight given conflicting haplotype *H*_1_and *H*_2_ as:

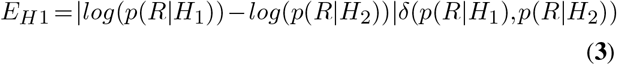

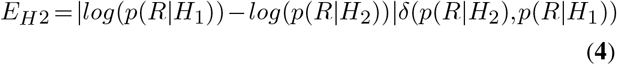

Where as *δ*(*a,b*) = 1 if *a*>*b* and *δ*(*a,b*) = 0 if *a*≤ b.

#### Interpretation of Fiedler vector

With the adjacency matrix of constructed linkage-graph, the haplotype is constructed based on the Fiedler vector. An expected Fiedler vector resembles the vector demonstrated in **Fig1:e**, in which elements corresponding to parallel alleles of given variant loci are assigned with opposite sign. The haplotype can then be constructed based on the element sign: alleles of different variant loci with the same sign belong to the same haplotype string. However, our experiments demonstrated that such Fiedler vector might not be generated occasionally. There are two exceptions where haplotype cannot be directly inferred from the Fiedler vector. The first case is trivial and it happens when the linkage graph contains pairs of variants loci that hold an equal score of two complement haplotypes. The resulted Fiedler vector possesses equal-signed elements of corresponding locus, which is pruned by our algorithm.

The second situation leads to a Fiedler vector that categorises variants into two sub-blocks, as demonstrated in **Fig1:g**. Such Fiedler vectors is commonly seen with error-prone TGS data and linked-reads where reads with the same barcode might originate from different DNA fragments. To infer haplotype with such Fiedler vector, we partition the adjacency matrix accordingly and apply spectral graph analysis respectively (see example in **Fig1:g**). This process is iteratively applied until SpecHap obtains a Fiedler vector which guides haplotype string. The sub-blocks are then merged together by treating each as a node and constructing a generalised linkage graph (**Fig1:h**)). The SpecHap algorithm might be summarised as below:

##### Algorithm 1: SpecHap haplotype assembling routine

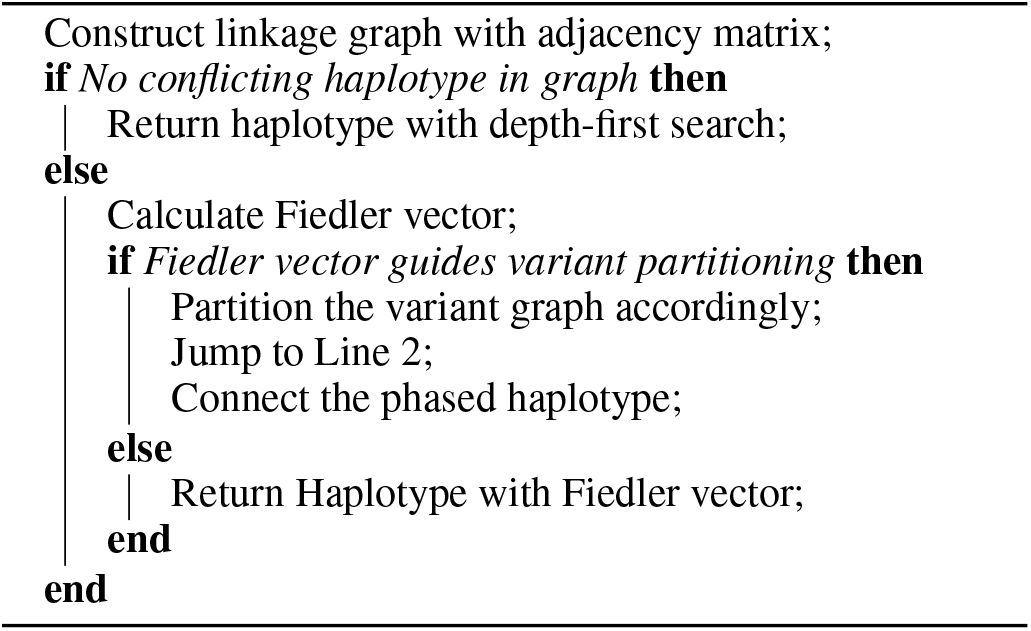

### Haplotype Assembling with Linked-Reads

In our experiments of phasing with 10x linked-reads, we identified major inconsistencies between the phased haplotype and the ground truth. Thus, special care was taken in order to obtain maximum phasing accuracy while preserving the completeness of assembled haplotype. First, variants are filtered by their allele depth and quality before the initiation of the phasing algorithm. We also disallow a phasing block from striding over 30 continuously filtered variants. Then, the covering range of each barcode is inferred based on the alignment results. In our implementation, a barcode can neither start nor end on an aligned read with mapping quality less than 30, and the overall barcode spanning length cannot be longer than 60kbp. This procedure reduces the rare but significant situations where reads with the same barcode are from two different DNA molecules.

### Haplotype Assembling with Hi-C

For Hi-C data that introduce transinteractions between homologous chromosomes, SpecHap treats possible transinteractions as general errors and does not model them specifically. SpecHap filters read pairs with insertion larger than 40M base-pair to avoid linkage with higher transinteration error rate.

### Chromosome Level Haplotype Construction

Whether all the heterozygous variants might be phased at the same time depends on the size and heterozygosity rate of genome. Since many mammalian species possess large genome, a divide and conquer strategy is applied. For simplicity in implementation, a definition of variant block and phasing batch is introduced. First, heterozygous variants that are supposed to be phased together may lie in one variant block. By default, the SpecHap assigns multiple blocks based on the input variant information. Blocks containing no sequencing noise could be phased directly by a graph depth-first search. Second, SpecHap could phase a batch, which maintains user-designated length, of the variant block at one time. Each pair of adjacent batches maintains an overlap of variant blocks, and the phased result of the previous batch is introduced when phasing its neighbor.

### Algorithm Complexity

The computation conducted by SpecHap comprises three aspects: depth-first search (DFS), calculation of graph Laplacian and Fiedler vector. The time complexity can be summarized as:

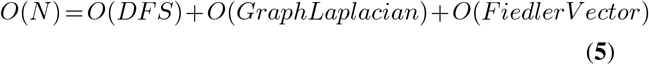

Define k as the number of eigenvector to be calculated, *n* = |*V* | (# of nodes in graph) and *m* = |*E* | (# of edges in graph). The time complexity of DFS is *O*(*n*+*m*). The computation of unnormalised graph Laplacian involves matrix addition and the complexity is *O*(*n*^2^). LU decomposition is adopted for eigen-calculation and the complexity to calculate k smallest eigenvectors is *O*(*kn*^2^). Therefore:

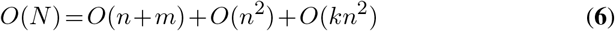

SpecHap calculates the first two eigenvectors to obtain the Fiedler vector, i.e. k = 2. Thus, the time complexity of the algorithm can be summarized as:

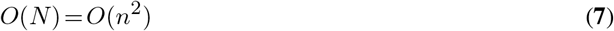

### Human Genome Dataset Processing

The phase3 trio-phased haplotype for NA12878 (aligned to hg19) was adopted as groud truth when benchmarking phasing accuracy.

*NGS data* The NGS alignment data for NA12878 were downloaded from the 1000 Genome Project.

*PacBio SMRT WGS data* PacBio SMRT WGS data were acquired from GIAB with NCBI accession number SRX1607993. Alignment was performed with minimap2 (Li, 2018) with preset parameters.

*Nanopore data* Nanopore reads were downloaded from ENA with accession PRJEB30620(Bowden *et al*., 2019). Alignment was performed with minimap2 (Li, 2018) with preset parameters.

*10x linked-reads data* 10x linked-reads for NA12878 were gathered from 10x genomics officials at https://support.10xgenomics.com/genome-exome/datasets/2.0.0/NA12878_WGS and alignment was performed with LongRanger2.2.1.

*Hi-C data* Hi-C sequencing data were downloaded from NCBI PRJNA473369. Sequenced reads from seven selected cells (SRR7226668, SRR7226671, SRR7226678, SRR7226679, SRR7226681, SRR7226682 and SRR7226685) were combined for further analysis. The alignment and insertion size of each read-pair were determined with BWA (Li and Durbin, 2009).

### *Ambystoma Mexicanum* Genome Dataset Processing

We also acquired the PacBio SMRT sequencing reads of an individual *Ambystoma mexicanum* with NCBI accession code PRJNA378970 (Nowoshilow *et al*., 2018). Alignment was performed by minimap2 to its chromosome-scale assembly GCA 002915635.2 (Smith *et al*., 2019). The called variant is accessible from EBI with accession number ERZ1668256.

### Sequencing Data Simulation

The trio-phased sample HG00403 were taken as the haplotype for simulation on chromosome 1, 21 and 22. Reads of length 150bp and insert size 350bp were simulated for NGS and linked-reads with wgsim and LRSIM (Luo *et al*., 2017) respectively with 30X coverage. 50X PacBio SMRT and ONT reads were also simulated based on same haplotype by PBSIM (Ono *et al*., 2012) and DeepSimulator (Li *et al*., 2018) with protocol-specific sequencing error rate.

## RESULTS

To address the problem of haplotype phasing in the era of TGS, linked-reads, and Hi-C reads, we developed SpecHap. Instead of assembling haplotype either by fragment partitioning or likelihood optimization, SpecHap utilises spectral graph theory that transforms the problem of haplotype phasing into an algebraic problem. SpecHap also incorporates protocol-specific filtering that improves phasing accuracy with Hi-C and 10x linked-reads. To access the efficiency and accuracy of SpecHap, we benchmarked its performance with existing methods on both simulated and WGS data. Previous publication (Duitama *et al*., 2012, Edge *et al*., 2017)s has already demonstrated the robustness of some state-of-art methods: FastHare (Panconesi and Sozio, 2004), RefHap (Duitama *et al*., 2010) and HapCUT2 (Edge *et al*., 2017). Some recent publications introduce novel methods, for instance, DCHap (Li and Lin, 2020), with improved efficiency and comparable accuracy. Therefore, we compared SpecHap with four existing methods: FastHare, RefHap, HapCUT2 and DCHap. All programs were benchmarked on CentOS with Intel Xeon CPU E7-4850 v2. CPU and peak memory usage were collected with Oracle Grid Engine. **Fig2** illustrates the overview of the conducted experiment setting and status. SpecHap managed to pass all experiment settings, while the other four software either not supported a specific data type or failed to complete the task within the usage boundary (24 CPU hours for *Homo sapien*, 5 CPU days and 40GB peak memory usage for *Ambystoma mexicanumm*).

**Figure 2.**
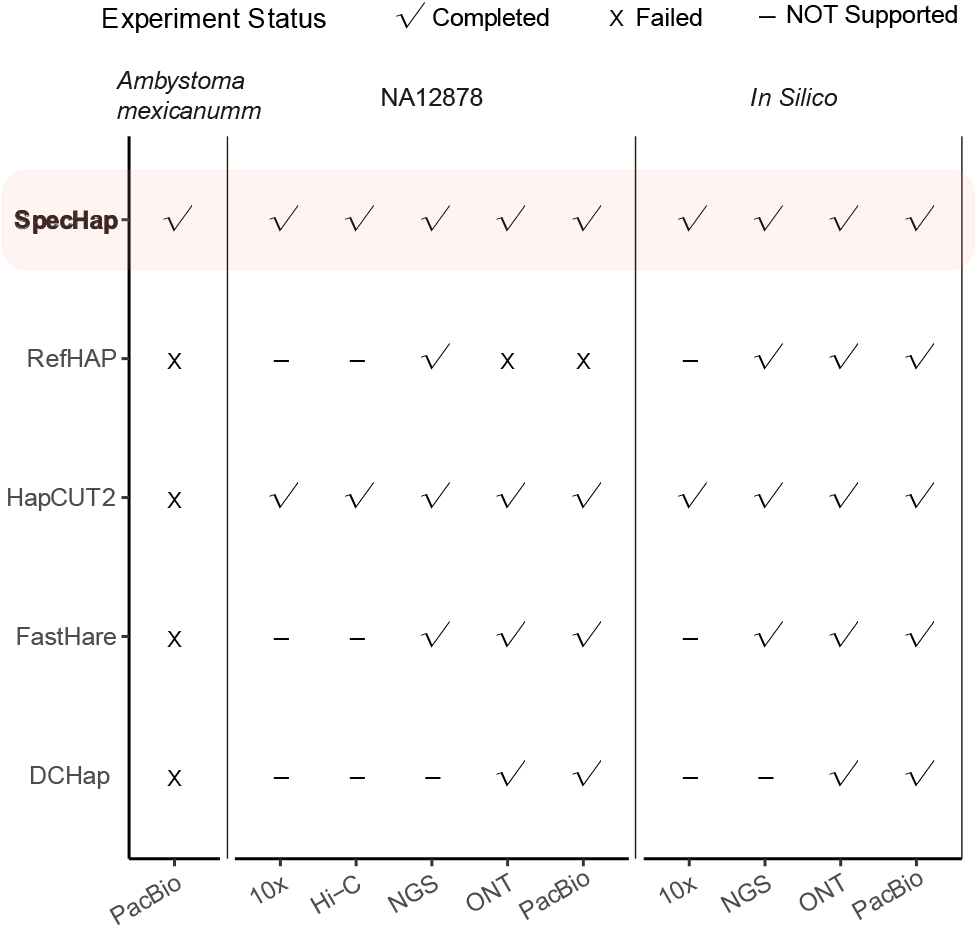
Overview of conducted experiment setting and status. Software that not support specific data type is marked with “-”. For *Homo sapien*, methods that exceed 24 CPU hours limits are marked as “Failed”. For *Ambystoma mexicanumm*, methods exceed 5 CPU days and 40GB peak memory usage are marked as “Failed”.

### SpecHap Demonstrated Runtime and Memory Efficiency

We first used *in silico* data to benchmark SpecHap with the existing method focusing on the third-generation sequencing protocol. Since most methods are not specifically designed for Hi-C and l0X Genomics linked-reads, these methods may not function as intended. We first simulated 50X PacBio SMRT reads and 50X ONT reads with PBSIM (Ono *et al*., 2012) and DeepSimulator (Li *et al*., 2018) based on the trio-phased haplotype for HG00403 from 1000 Genomes Project on chromosome 1, 21 and 22. To ensure a fair comparison, all the methods adopted the same set of fragments as input and default parameters are used for all sequencing protocol and methods. The CPU time and peak memory usage for all the methods were summarised in **Fig3:a** and Supplementary Table S1.

**Figure 3.**
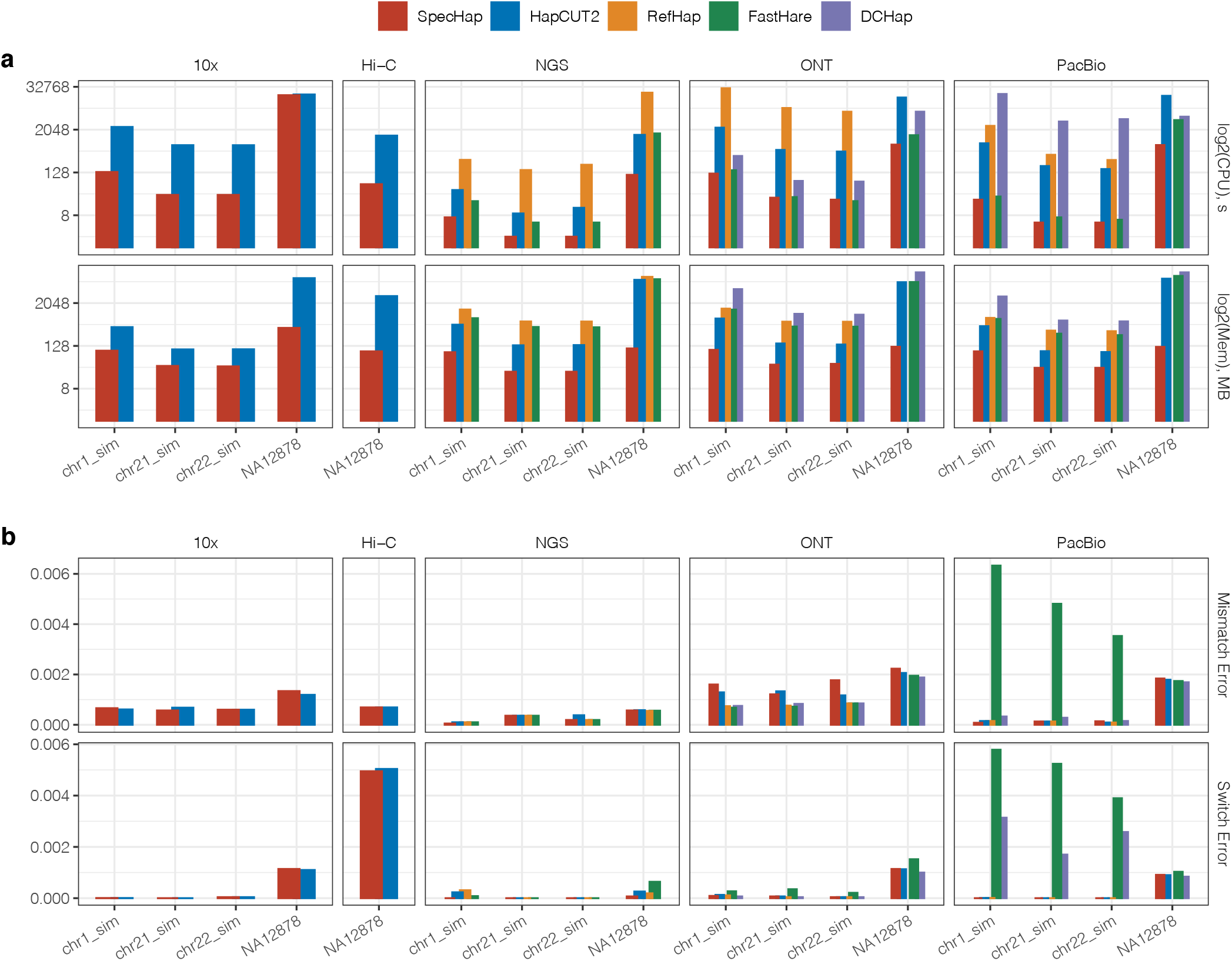
Overall benchmarked data on both simulation and WGS sample NA12878 for diverse sequencing protocol. **a** Log2-scaled CPU and peak memory usage, in second and MegaBytes respectively. Value exceeding 30,000 CPU seconds and 15GB peak memory usage is capped. **b** Overall switch error rate and mismatch error rate, calculated as the number of error divided by the number of possible error.

When phasing with PacBio SMRT sequence, SpecHap outperformed all four existing methods considering both CPU time and peak memory usage based on the simulation. SpecHap was around 40 times faster than HapCUT2, 100 times faster than RefHAP and as fast as FastHare. DCHap did not function efficiently and accurately enough on simulations with PacBio SMRT reads. While assembling haplotype with ONT reads, SpecHap still achieved 20 times faster than HapCUT2, 30 times faster than RefHAP, 2 times faster than DCHAP and as fast as FastHare. FastHare, the second most efficient methods, demonstrated significant higher switch error rate while comparing with other methods (**Fig3:b** and Supplementary Table S2) on both PacBio SMRT and ONT sequence. The runtimes and memory comparison of NGS, Hi-C and 10x linked-reads can be found in Supplementary Table S1:a-c.

As for WGS sample NA12878, SpecHap persisted with its efficiency. It is also worth mentioning that SpecHap consumed minimum peak memory (Mega-Bytes level) compared with other methods (Giga-Bytes level) on WGS sample. With WGS 10x linked-reads, SpecHap and HapCUT2 achieved comparable speed. However, HapCUT2 requires additional computation on fragment linking (32 CPU hours on the linkage of fragment with the script provided by HapCUT2).

### SpecHap Accurately Phased Human Individual NA12878 on Diverse Sequencing Protocols

We assessed the accuracy of SpecHap with an individual NA12878 with five sequencing protocols: NGS, 10x Genomics linked-read, Hi-C, PacBio SMRT and ONT sequencing. The trio-phased haplotype of NA12878 from the 1000 Genomes Project was taken as the ground truth and input to assemble haplotype for each data type. The measurement was conducted based on the metrics of mismatch error rate and switch error rate (**Fig3:b** and **Fig4:a-b**), which are widely adopted criteria while accessing the accuracy of the assembled haplotype (Edge *et al*., 2017). As for the completeness of the assembled haplotype, AN50, which is the N50 of adjusted haplotype span such that half of all phased heterozygous variant loci are in the segment, and the number of phased SNVs were adopted (**Fig4:c-d**, Supplementary Table S3).

**Figure 4.**
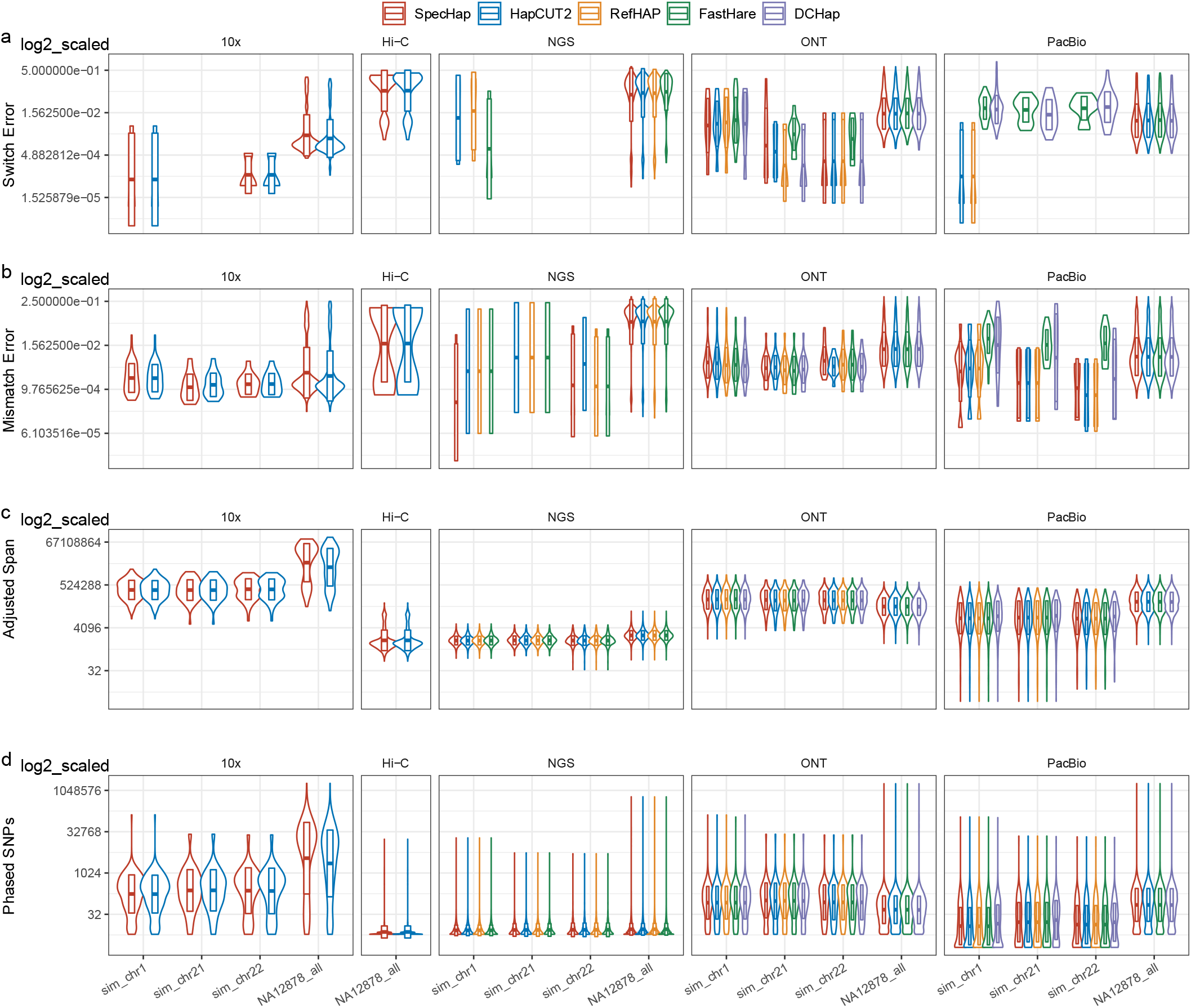
Log2 scaled violin plot of benchmarked data on per-phased-block scale. **a** Switch error rate. Methods with zero switch error on simulation dataset are not showed (10x chr21 sim, NGS chr21 sim, chr22 sim, PacBio chr1 sim, chr21 sim and chr22 sim). **b** Mismatch error rate. **c** Adjusted span, defined as phased block span times ratio of number of phased SNPs over number of total SNPs, in base pair. **d** Number of phased SNPs.

#### NGS data

To access SpecHap on NGS data, we took a high coverage PCR-free sequenced data for individual NA12878 from the 1000 Genomes Project. As displayed in **Fig3:a**, the switch error rate for SpecHap was less than RefHap, the second most accurate algorithm among all other software. HapCUT2 shared virtually similar switch error of RefHap while keeping more variants unpruned. FastHare, however, demonstrated higher switch error rate with more pruned record comparing with SpecHap and HapCUT2. As for the mismatch error rate, all four methods shared comparable results with no statistical difference (**Fig3:b** and Supplementary Table S2-a). SpecHap completed the haplotype assembling with the least CPU (2 CPU minutes) and peak memory (109MB) while preserving the phasing accuracy (Supplementary Table S1-a).

#### Hi-C data

Since most methods are not specialized for haplotype assembling with Hi-C, we only compared the quality of haplotype assembly generated by SpecHap and HapCUT2. HapCUT2 models the translocation error by iteratively estimating the probability of reads originating from different homologous chromosomes(Edge *et al*., 2017). However, SpecHap demonstrated lower comparable switch error rate on the specific data set without modeling translocation error (**Fig3:b** and Supplementary Table S2-c). SpecHap also outperformed HapCUT2 on computational load by completing the haplotype assembly within 1 CPU minute (Supplementary Table S1-c).

#### 10x linked-read data

We benchmarked SpecHap on 10x genomics linked-reads. The 10x linked-reads label short reads that originated from a single long DNA fragment with the same barcode. Although DNA fragment generally spans around 60k base-pair range, reads might be sparsely distributed across the original fragment. It is also possible that two reads who shared the same barcode originated from different DNA molecules. The data set we acquired has around 50X coverage and reads were filtered that fragments with white-listed barcode were kept. The same set of variants was used to extract fragment information. SpecHap achieved a comparable accuracy by completing the assembly with 5 CPU hours while HapCUT2 spent 5.6 CPU hours (Supplementary Table S1-b). Moreover, SpecHap takes raw fragment as input while other methods requires extra computation on fragment linking. The linkage of fragment was conducted with the script provided by HapCUT2 and took 32 CPU hours to complete.

#### PacBio SMRT data and ONT data

PacBio SMRT sequencing and ONT sequencing are known for their error-prone (∼10%) long reads. We compared the performance of SpecHap with other methods on 44X coverage PacBio reads and 40X coverage ONT reads. Fragment information was extracted with the same set of variants call. On PacBio SMRT data, SpecHap achieved the fastest speed (21 CPU minutes) while achieving virtually identical switch error rate comparing with HapCUT2 (**Fig3:a** and Supplementary Table S1-d). FastHare, with second-best efficiency, demonstrated higher switch error rate with most variant pruned (**Fig3:b**). DCHap, with slightly more accurate result, assembled the haplotype with the lowest AN50 and pruned the second most variants (Supplementary Table S3-d). All methods demonstrated no significant statistical differences considering the mismatch error rate. On ONT data (**Fig3:a** and Supplementary Table S1-e), SpecHap persisted its efficiency by completing the assembly within 17 CPU minutes with comparable switch error rate. DCHap, with the least switch errors, pruned more than 2000 SNVs than SpecHap and HAPCUT2 (Supplementary Table S3-e). RefHap failed to finish the assembly process with both PacBio SMRT reads and ONT reads due to excessive time consumption and accuracy estimation was not available.

### SpecHap Demonstrated Scalability by Phasing with Large Diploid Genome

SpecHap also demonstrated the scalability by assembling the 32X PacBio SMRT sequenced reads with N50 read length around 14.2kbp of an *Ambystoma mexicanum* individual which posses 32 billion base-pair-long genome (Nowoshilow *et al*., 2018, Smith *et al*., 2019). The variant set of the same individual that containing more than 20 million heterozygous SNV loci. As illustrated in Supplementary Figure 1, SpecHap were able to finish the assembly within 6 CPU hours with only 945MB of peak memory consumption, while all other methods failed to finish within time (5 CPU days) and memory limit (40GB).

## DISCUSSION

SpecHap, a novel diploid phasing algorithm, supports sequencing data with different coverage from diverse platforms. SpecHap transforms haplotype phasing into a linear algebra problem by applying spectral graph theory. The Fiedler Vector guided allele partitioning might be interpreted as min-cut on our linkage graph (Von Luxburg, 2007). To our knowledge, this is the first time to employ spectral graph theory in haplotype phasing. Although there is no guarantee that SpecHap provides optimal solutions, we demonstrated that our model succeeded in efficiently and accurately assembling haplotypes with diverse sequencing reads from individual NA12878. SpecHap is also scalable to phase one of the largest diploid genomes of *Ambystoma mexicanumm*.

Long-reads sequencing introduces significant advances considering the completeness of assembled haplotype. However, SMRT and ONT long-reads maintain higher error rate per base and may fail to accurately detect SNVs, particularly heterozygous ones (Edge and Bansal, 2019). In our experiments, we adopted the trio-phased high-quality variants set for NA12878 from the 1000 Genome Project as input for all methods. Since most SIH phasing methods including SpecHap requires a high-quality set of variants, 30X Illumina short reads sequencing might be performed to obtain reliable calls for SNVs and INDELs. Some recent approaches based on deep learning were also able to identify variants accurately from long-reads sequencing (Luo *et al*., 2019).

In addition to the wet-lab based phasing (Redin *et al*., 2019), computational approaches including population-based phasing, Mendel’s Law based phasing and SIH are also widely adopted (Majidian and Sedlazeck, 2020). **(i)** Although performed at a lower coverage, the population based phasing requires a haplotype population structure model and can only phase common variations (Lutgen *et al*., 2020). Genomes of multiple non-human individuals in one population under natural condition are often difficult to obtain. **(ii)** A chromosome-scale haplotype might be constructed by applying the Mendelian laws of inheritance (also known as trio-based phasing) (Blackburn *et al*., 2020). However, it lacks the ability to phase *de novo* mutations, especially in Mendelian diseases. Trio-based phasing usually requires high coverage sequencing and sequences from both parents (Delaneau *et al*., 2012, Garg *et al*., 2016). **(iii)** Compared with previously mentioned phasing methods, SIH is the most comprehensive approach as it incorporates de novo mutations and rare variations. Although NGS usually brings the possibility of hindering due to fragmentation, the development of linked-reads and TGS long-reads introduces significant advance on phasing quality of SIH (Browning and Browning, 2011, Sedlazeck *et al*., 2018). Thus, we developed SpecHap here, a novel method for haplotype assembly compatible with diverse sequencing protocol.

Although WGS phasing has grown as an easy-to-handle approach for studying genetic variations within or between species with “normal” genome size, this technology is immature for non-model large genomes yet (Weisrock *et al*., 2018). From the perspective of conservation, larger genomes are assumed to become particularly vulnerable during periods of increased extinction (Vinogradov, 2003). Also, lineages with giant genomes seem to be eliminated from the most extreme habitats (KNIGHT *et al*., 2005). The largest genomes are presently unearthed in amphibian salamanders, with sizes ranging from 14-120 gigabytes (Newman and Austin, 2016). Salamanders, the non-model species, represent the most massive regenerative repertoire, capable of rebuilding and restoring multiple organs and systems such as brain structures, and has the potential to become model organisms (Elewa *et al*., 2017). The genome of Mexican salamander, axolotl, with a size (∼32 gigabytes) approximately ten times bigger than homo sapiens’, is one of the largest genomes (Nowoshilow *et al*., 2018). Due to the high raising and sequencing cost for a cohort of axolotls, scientists virtually rely on an individual axolotl genome assembly other than population-based phasing to conduct biological investigations (Sotero-Caio *et al*., 2017). Nevertheless, our experiments illustrated that some state-of-art haplotype assembly tools failed to handle the oversized genomes. We managed to phase the haplotype of axolotl with PacBio SMRT sequencing data, implying SpecHap might be a compatible tool for future research on the evolutionary history of amphibians and other organisms with immense genome scale.

In this study, we introduced a novel diploid SIH phasing algorithm SpecHap and demonstrated its robustness and efficiency. By transforming haplotype phasing into a linear algebra problem, SpecHap assembled haplotype with ultra-fast speed while preserving comparable accuracy. Moreover, a comprehensive analysis on the influence of technological specific error over phasing quality may be conducted for Hi-C and 10x linked-reads. Although our algorithm works on diploid genome, generalization towards high ploidy is expected. Besides, determination of haplotype-resolved structural variation might also be an important feature to be introduced in the future.

## ACKNOWLEDGEMENTS

We would like to express sincere gratitude to Dr. Yen Kaow Ng from Kotai Biotechnologies, Japan for manuscript revision and to Dr. Zijun Xiong of the Chinese Academy of Sciences for suggestions on data collection. We appreciate Miss Santu for her creation of an image for *Ambystoma mexicanumm*. We would also like to thank Dr. Wenlong Jia and Mr. Bowen Tan for their valuable assiatance and advice.

## Conflict of interest statement

None declared.

### Data Availability

All the data used in this paper can be retrieved from public database. All the experiments are reproducible with dedicated version of software with default arguments.

### Software Availability

SpecHap source code is deployed at https://github.com/deepomicslab/SpecHap and a copy is attached in the supplemental material. The fragment information was extracted with a modified version of extractHAIRS (Edge *et al*., 2017) which is packed with SpecHap. We adopted the implementation of FastHare from Duitama *et al*. (2012) for benchmarking.

## Notes

### Competing Interest Statement

The authors have declared no competing interest.

